# Accuracies of univariate and multivariate genomic prediction models in African Cassava

**DOI:** 10.1101/116301

**Authors:** Uche Godfrey Okeke, Deniz Akdemir, Ismail Rabbi, Peter Kulakow, Jean-Luc Jannink

## Abstract

**List of abbreviations:**

GS: Genomic Selection
BLUP: Best Linear Unbiased Prediction
EBVs: Estimated Breeding Values
EGVs: Estimated genetic Values
GEBVs: Genomic Estimated Breeding Values
SNPs: Single Nucleotide polymorphisms
GxE: Genotype-by-environment interactions
GxE: Genotype-by-environment interactions
GxG: Gene-by-gene interactions
GxGxE: Gene-by-gene-by-environment interactions
uT: Univariate single environment one-step model
uE: Univariate multi environment one-step model
MT: Multi-trait single environment one-step model
ME: Multivariate single trait multi environment model

**Abstract:** *Background:* Genomic selection (GS) promises to accelerate genetic gain in plant breeding programs especially for long cycle crops like cassava. To practically implement GS in cassava breeding, it is useful to evaluate different GS models and to develop suitable models for an optimized breeding pipeline.

*Methods:* We compared prediction accuracies from a single-trait (uT) and a multi-trait (MT) mixed model for single environment genetic evaluation (Scenario 1) while for multi-environment evaluation accounting for genotype-by-environment interaction (Scenario 2) we compared accuracies from a univariate (uE) and a multivariate (ME) multi-environment mixed model. We used sixteen years of data for six target cassava traits for these analyses. All models for Scenario 1 and Scenario 2 were based on the one-step approach. A 5-fold cross validation scheme with 10-repeat cycles were used to assess model prediction accuracies.

*Results:* In Scenario 1, the MT models had higher prediction accuracies than the uT models for most traits and locations analyzed amounting to 32 percent better prediction accuracy on average. However for Scenario 2, we observed that the ME model had on average (across all locations and traits) 12 percent better predictive power than the uE model.

*Conclusion:* We recommend the use of multivariate mixed models (MT and ME) for cassava genetic evaluation. These models may be useful for other plant species.

## Background

Cassava (*Manihot esculenta* Crantz) [1] is a staple food for over 700 million people in Africa, South America and Asia [2], Cassava also has immense industrial potential. Pure white cassava starch is easy to extract and contains low levels of fat (about 1.5%), protein (about 0.6%) and phosphorus (about 4%), which are desirable attributes for the food industry [3,4], Given the issues of climate change and rapid population growth in countries that rely heavily on cassava, rapid genetic improvement of cassava is critically needed. To enable rapid genetic improvement of cassava, genetic evaluation protocols hinged on Best Linear Unbiased Prediction (BLUP) analysis [5,6] and selection on a merit index [7,8] has been recommended [9] to maximize gain from selection.

Genomic selection (GS) [10] offers crops like cassava tremendous opportunity for accelerated genetic gains [11] by making use of whole genome SNP markers accessible with methods like the genotyping-by-sequencing (GBS) [12]. These whole genome SNP markers could be dense enough to be in linkage disequilibrium with most quantitative trait loci (QTL) affecting traits of interest. Using GS, selection is imposed at these QTL without actually identifying the QTL or the functional polymorphisms [10]. Also these markers will help to better track relatedness due to mendelian sampling [24] in a breeding population especially where pedigree records are not fully available thus yielding an improvement in selection accuracies [25].

## GS models for plant genetic evaluation

Genetic evaluation [9] starts with accurately estimating the genetic value of an individual for a wide range of traits using its own performance records, progeny performance records or records from relatives or a combination of the three [13]. This has usually been carried out using single trait (uT) BLUP methodology [14] for obtaining estimated breeding values (EBVs) for one trait at a time. In plant and animal breeding, breeders usually select on the basis of multiple traits that are often genetically correlated although these correlations may range from very weak to very strong. The uT model for traits measured in a single environment assumes zero genetic and residual covariances between these traits such that information from other traits are not utilized when obtaining EBVs of the evaluated individuals for the traits in the analysis. However, the optimal estimation procedure to combine information from multiple trait records and obtain EBVs is the multi-trait BLUP methodology (MT) [15,16]. The MT model does not assume zero genetic and residual covariances but rather provides an estimate for these and also uses this information when obtaining individual EBVs for the traits in the analysis. The MT model has several advantages over the uT model including:

▪ Improved prediction accuracies for individual traits in the model because of more information (direct or indirect), better connectedness of the data [17], and exploitation of genetic and residual correlations in the model especially when traits with varying heritabilities are analyzed jointly.
▪ Simplified index selection because optimal weight factors for the total merit index are the economic weights [17].
▪ Efficient procedures for obtaining genetic and residual covariances and EBVs across location, country or region evaluations on information from same or related individuals [18, 19].
▪ Better selection accuracies by accounting for selection bias when all target traits under selection are included in the model [20] in addition to the use of all individuals (selected or not) in the relationship matrix.

In most plant breeding programs, selected individuals are evaluated in multi-environment trials (METs). The goal is to sample the influence on selection candidates of the range of environments for which varieties will be targeted. Addressing the problem of the analysis of METs brings into focus another potential use for MT models [30]. Here, phenotypes of the same trait, but measured at different locations are parameterized as different traits in the MT model [31], producing what we call a multi-environment BLUP (ME) model. Like the MT model, the ME model estimates genetic covariances between a single trait measured at multiple environments which may lead to more accurate estimates of individual EBVs for the trait at all the environments where data has been recorded. For ME models used for modelling MET data, residual covariances are set to zero reflecting the assumption that no mechanism generates error covariances between a trait measured in different environments [18]. In contrast, the typical univariate BLUP model for modelling METs data, termed the univariate multi-environment model (uE), fits a multikernel mixed model with the genotypic effect as one kernel and the genotype-by-environment (GxE) effect as the second kernel and maybe environment as third kernel [26]. This model yields a GxE variance for a MET and individuals can be ranked on their performance at different locations. It is expected that for traits with high correlation between environments, the uE model should offer more accurate EBVs, while EBV accuracies for traits with lower correlation between environments will benefit more from the ME model. Different variants of the ME model have been used for modeling environment covariance structures in plant [32-35] and in animal breeding [36,37]. Genetic covariances from the ME model offer a convenient tool for assessing the impact of GxE on a trait. The genetic covariances relate directly to the extent of GxE at all locations in the analysis. A low genetic correlation of the EBVs between a trait at different locations from the ME model indicate GxE impact on that trait [9,38-41].

Selecting the GS model to be employed in a practical cassava breeding program is not trivial hence the need to compare different GS models that will be useful in the different stages of cassava breeding with METs data. Finally, fitting multivariate BLUP models is not trivial. Even with software that can in principle fit these models, model convergence is not guaranteed and may require several attempts [21-23] and hence univariate models may be more practical if benefits of the multivariate models are not substantial.

The objectives of this paper are to:

▪ Compare multi-trait (MT) and single trait (uT) mixed models for METs data using cross-validated prediction accuracies.
▪ Compare the multivariate multi-environment (ME) model to a single-trait multi-environment (uE) model using cross-validated prediction accuracies while assessing GxE impact on analyzed traits via genetic covariances from the ME model fit.

## METHODS AND MATERIALS

### Cassava phenotype data

We used historical phenotype data from different trials conducted by the cassava breeding program at the Institute of Tropical Agriculture (IITA), Ibadan, Nigeria in our analysis. The Genetic Gain population represents a collection of clones selected from the 1970s to 2007 by the cassava breeding program at the IITA [48,49]. Some of these clones are West African landraces and some are of East African origin. Clones in the Genetic Gain population have gone through advanced stages of the cassava breeding process up to on farm variety testing trials. The data used in our analysis comprises data collected on clonal evaluation trials (CETs) which are augmented design trials with about 2 known checks and unreplicated plots with 5 plant stands per clone. These data were collected from three target locations in Nigeria: Ibadan (7.40° N, 3.90° E), Mokwa (9.3° N, 5.0° E), and Ubiaja (6.66° N, 6.38° E). These locations represent regions which encompass about 35 percent of the cassava production base in Nigeria. Data sets were collected from 2000 to 2015 and included trials with most of the 764 clones of the Genetic Gain population. Six target agronomic traits were used in the analysis including seedling vigor (VIGOR), Number of storage roots per hectare at harvest (RTNO), Fresh weight of harvested roots expressed in tons per hectares (T/ha) per plant (FYLD), Percentage dry matter (DM) of storage roots, which measures root dry weight as the percentage of the root fresh weight, plot mean cassava mosaic disease severity (MCMDS), rated on a scale from 1 (no symptoms) to 5 (extremely severe), and plot mean cassava green mite (MCGM) severity, rated on a scale from 1 (no symptoms) to 5 (extremely severe). Cassava mosaic disease is caused by a *Begomovirus* that belongs to the Geminiviridae family, and is carried and transmitted by the whitefly *Bemisia tabaci.* The cassava green mite is *Mononychellus tanajoa* [50]. These traits are target traits used in the selection index for selection decisions in the IITA cassava breeding program. Phenotype data metrics are shown in Table 1.

**Table 1.** Cassava phenotype data metrics showing number of records, number of observed clones and mean of traits for 3 locations Ubiaja, Mokwa and Ibadan.

|  | **Ubiaja** | **Mokwa** | **Ibadan** |
| --- | --- | --- | --- |
| **No. of records** | 7806 | 5345 | 6010 |
| **No. of Clones** | 739 | 573 | 691 |
| **VIGOR** | 6.51 | 6.52 | 6.11 |
| **RTNO** | 31.71 | 64.05 | 52.74 |
| **FYLD** | 12.61 | 16.51 | 15.84 |
| **DM** | 31.95 | 29.01 | 35.8 |
| **MCMDS** | 1.59 | 1.21 | 1.93 |
| **MCGM** | 3.56 | 2.99 | 3 |

### Cassava genotype data

DNA was extracted using DNeasy Plant Mini Kits (Qiagen) from 739 clones from the 2013 Genetic Gain trial at IITA and was quantified using PicoGreen. Genotyping-by-sequencing (GBS) was used for genotyping [12] these clones. Six 95-plex and one 75-plex ApeKI libraries were constructed and sequenced on Illumina HiSeq, one lane per library. Single nucleotide polymorphisms (SNPs) were called from the sequence data using the TASSEL pipeline version 4.0 [51], using an alignment to the *Manihot esculenta* version 6 reference genome [52]. Individuals with greater than 80% missing SNP calls and markers with more than 60% missing were removed. The marker data was converted to dosage format (0, 1, 2) and missing genotypic data were imputed using a LASSO regression method (Ariel Chan, personal communication, 2014) implemented using the R glmnet package [53]. The final data set consisted of 183,201 SNPs scored in 764 clones.

### Statistical analysis

We restructured the cassava phenotype data described above into two types of data common in most plant breeding programs. The first set was achieved by pooling data from multiple years in all locations (multi-environment trials data - METs). We termed this scenario the single environment genetic evaluation (Scenario 1). This resulting predictive power from this data were assessed for our three locations Ubiaja, Mokwa and Ibadan. The second scenario was achieved by using same METs data but in this case extracting location specific information by attempting to model GxE interaction. We termed this scenario the multi-environment genetic evaluation (Scenario 2). The goal of the latter data is to access the value of evaluate the impact of GxE and check if this yields better predictive value of the breeding value of a clone.

### Pseudo-true genetic values for model accuracy computations

For validating the models in this study, we define first a univariate single trait mixed model for each trait at each location separately (to preserve the variation embedded in each location) using an identity covariance matrix among clone effects, which assumes no relationship among all clones. The univariate mixed model was as follows:

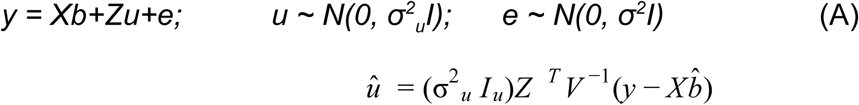

where *y* is a vector of observations, *b* is a vector of fixed effects with design matrix *X* (relating observations to fixed effects in this case including grand mean and a nested effect of Trial-within-Year-within-Location and the ratio of plants harvested to number planted); *u* is a vector of clonal genetic effects with design matrix *Z* (relating observations to clones). This model was fit using the *Imer* function in the R Ime4 package [55] and resulting BLUP values (*û*), which we refer to as Estimated Genotypic Values (EGVs), were used as pseudo-true genetic effects for prediction accuracy computations.

### GS models for Scenario 1

We define two mixed models fitted here as follows:

#### The single trait mixed model (uT)

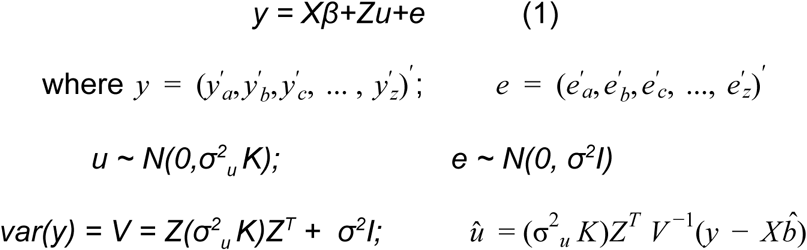

where *y* is the response vector of a trait at locations *a, b.. z, β* is the vector of fixed effects with design matrix *X* (relating observations to fixed effects namely the grand mean, nesting of Trial-within-Year-within-Location and ratio of plants harvested to number planted); *u* is the vector of random additive genomic effects with design matrix *Z* (relating trait values to clones) and *K* is the additive genomic relationship matrix generated from SNP markers as in VanRaden, 2008 [63] method two. Estimation of the parameters in model (1) were performed using Restricted Maximum Likelihood (REML) procedure implemented in the *airemlf90* FORTRAN program of the *blupf90* package [62] which estimates (co)variance parameters and then obtains best linear unbiased estimates (BLUEs) of fixed effects and BLUPs of random effects by solving the mixed model equations (MME) [5,6].

#### The multitrait Mixed Model

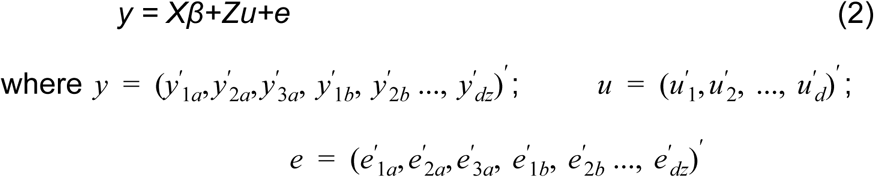

where *y* is a long vector of *d* traits in locations *a, b … z*, recorded for *n* clones, **X** and **Z** design matrices are block diagonal matrices represented as *diag* (X_a_, X_b_,…, X_z_) and *diag* (Z_a_, Z_b_,…, Z_z_) respectively allowing for missing clones and observations. **X** is a design matrix for fixed effects *β* (with components as in model 1 above for every location and trait) and **Z** is a design matrix for random genetic effects *u*. Following a multivariate normal distribution (*N_m_*), the marginal density of *y* is given as:

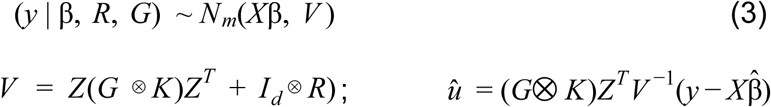

where *d* is number of traits being analyzed, G and R are *d* × *d* symmetric genomic and error covariance matrices respectively, *K* remains the additive genomic relationship matrix for *n* clones generated from SNP markers as in VanRaden, 2008 [63] method two, *I* is a *d*-traits identity matrix and u are the genomic estimated breeding values (GEBVs) the clones and for the traits in the analysis. Estimation of the parameters in model (3) were also performed using REML procedure implemented in the *airemlf90* program [62] from which BLUEs of fixed effects and BLUPs of random effects were also obtained by solving the mixed model equations (MME) [5,6].

### GS models for Scenario 2

We also defined two mixed models here with the aim of modeling genotype-by-environment interaction effects and these are as follows:

#### The univariate multi-environment model (uE)

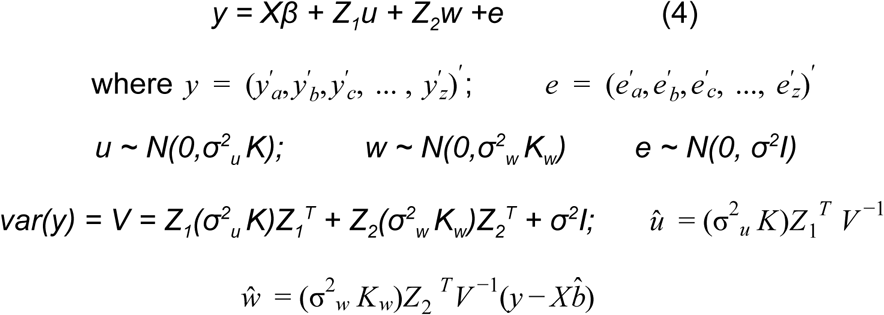

where *y* is a long vector of trait values for all locations, *β* is the vector of fixed effects with design matrix *X* (relating observations to fixed effects namely the grand mean, nested Trial-within-Year-within-Location and ratio of plants harvested to number planted); *u* is the vector of random additive genomic effects with design matrix *Z*_1_ (relating trait values to clones), *w* is the vector of random marker-based clone-by-location interaction effects with design matrix Z_2_ (relating trait values to clones-location combinations) and *K* is the additive genomic relationship matrix generated from SNP markers as in VanRaden, 2008 [63] method two, *K_w_* is a block diagonal covariance matrix represented as *diag(K*_a_*, K_b_, K_c_,…, K_z_*) with the *K* matrices in this block being genomic relationship matrices calculated as in VanRaden, 2008 [63] method two for clones in locations *a,b…z.* In this model, the genomic value of a clone were estimated as *û* + *ŵ.* We also fit model(4) using the *airemlf90* tool of the *blupf90* package [62] with a custom R-script for a 5-fold cross validation procedure with 10 repeat runs using the *airemlf90* function.

#### The multivariate multi-environment (ME) model

We fit the ME model in a single step procedure using the following model:

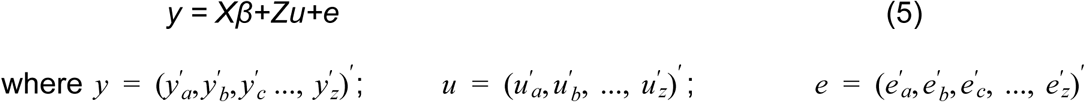

where *y* is a long vector of same trait in locations *a, b … z,* recorded for *n* clones, **X** and **Z** design matrices are block diagonal matrices represented as *diag*(X_a_, X_b_,…, X_z_) and *diag* (Z_a_, Z_b_,…, Z_z_) respectively allowing for missing clones and observations. **X** is a design matrix for fixed effects *β* (with components as the grand mean and year effect) and **Z** is a design matrix for random genomic effects *u.* Following a multivariate normal distribution *(N_m_)*, the marginal density of y is given as:

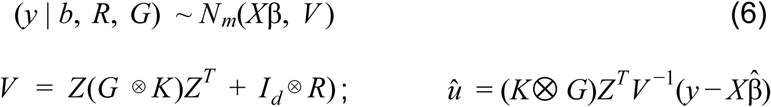

where *d* is number of traits being analyzed, G and R are *d* × *d* symmetric genomic and error covariance matrices respectively, *K* remains the additive genomic relationship matrix for *n* clones generated from SNP markers as in VanRaden, 2008 [63] method two, *I* is a *d*-traits identity matrix and *u* are the genomic estimated breeding values (GEBVs) the clones and for the traits in the analysis. Estimation of the parameters in model (6) were also performed using REML procedure implemented in the *airemlf90* program [62] from which BLUEs of fixed effects and BLUPs of random effects were also obtained by solving the mixed model equations (MME) [5,6]. In this model, the error covariance matrix R is a diagonal matrix allowing heterogeneous variances of a trait for different locations but the covariances are restricted to zero following the assumption that no mechanism generates errors between a trait at multiple locations.

### Comparison of prediction accuracies

We used a 5-fold cross validation scheme with 10 repeats for comparisons between the univariate and and multivariate models. The same folds were used for the models in each scenario. We hereafter refer to predicted BLUPs or genomic effects from these models as Genomic EBVs (GEBVs). Prediction accuracies were calculated as a correlation of the validation fold GEBVs to their corresponding EGVs.

## Results

### Scenario 1: MT vs uT models

In Scenario 1, we observed that the prediction accuracies of the MT model were higher than those from the uT models for most traits and locations in our analysis (Table 2). On average (across traits and locations), the MT model had 35 percent better predictive power for VIGOR, 34 percent for RTNO, 19 percent for DM, 33 percent for MCMDS, 54 percent for FYLD and 15 percent for MCGM compared to the uT model. The MT models had 32 percent higher predictive power than the uT models for all traits and locations in the model.

**Table 2.** Cross validation prediction accuracies for GS models in scenarios 1 and 2. Table 2 shows prediction accuracies for MT and uT models (GS scenario 1) and for ME and uE models (GS scenario 2).

| GS Scenario 1 |  |  |  |  |  |  |
| --- | --- | --- | --- | --- | --- | --- |
| Single trait single env. model (uT) |  |  |  | Multi-trait models (MT) |  |  |
|  | Ubiaja | Mokwa | Ibadan | Ubiaja | Mokwa | Ibadan |
| **VIGOR** | 0.25 ± 0.01 | 0.11 ± 0.01 | 0.38 ± 0.02 | 0.39 ± 0.02 | 0.09 ± 0.03 | 0.52 ± 0.03 |
| **RTNO** | 0.27 ± 0.01 | 0.19 ± 0.01 | 0.31 ± 0.01 | 0.39 ± 0.02 | 0.21 ± 0.03 | 0.43 ± 0.02 |
| **DM** | 0.59 ± 0.01 | 0.33 ± 0.01 | 0.49 ± 0.01 | 0.69 ± 0.01 | 0.39 ± 0.02 | 0.60 ± 0.02 |
| **MCMDs** | 0.49 ± 0.01 | 0.43 ± 0.01 | 0.56 ± 0.02 | 0.68 ± 0.02 | 0.58 ± 0.04 | 0.71 ± 0.01 |
| **FYLD** | 0.32 ± 0.01 | 0.13 ± 0.02 | 0.29 ± 0.01 | 0.46 ± 0.01 | 0.26 ± 0.03 | 0.42 ± 0.03 |
| **MCGM** | 0.36 ± 0.01 | 0.50 ± 0.01 | 0.56 ± 0.01 | 0.44 ± 0.02 | 0.54 ± 0.01 | 0.65 ± 0.01 |
| GS Scenario 2 |  |  |  |  |  |  |
| Single trait multi-env. model (uE) |  |  |  | Multi-env model (ME) |  |  |
|  | Ubiaja | Mokwa | Ibadan | Ubiaja | Mokwa | Ibadan |
| **VIGOR** | 0.22 ± 0.01 | 0.10 ± 0.01 | 0.37 ± 0.01 | 0.24 ± 0.01 | 0.12 ± 0.02 | 0.32 ± 0.01 |
| **RTNO** | 0.29 ± 0.01 | 0.11 ± 0.01 | 0.34 ± 0.01 | 0.27 ± 0.02 | 0.13 ± 0.01 | 0.34 ± 0.02 |
| **DM** | 0.49 ± 0.01 | 0.20 ± 0.02 | 0.40 ± 0.01 | 0.60 ± 0.01 | 0.35 ± 0.01 | 0.50 ± 0.01 |
| **MCMDs** | 0.40 ± 0.01 | 0.23 ± 0.01 | 0.53 ± 0.01 | 0.48 ± 0.01 | 0.39 ± 0.02 | 0.57 ± 0.01 |
| **FYLD** | 0.38 ± 0.01 | 0.10 ± 0.02 | 0.35 ± 0.01 | 0.37 ± 0.01 | 0.12 ± 0.03 | 0.36 ± 0.02 |
| **MCGM** | 0.31 ± 0.01 | 0.48 ± 0.01 | 0.56 ± 0.01 | 0.38 ± 0.02 | 0.47 ± 0.01 | 0.55 ± 0.01 |

### Scenario 2: ME vs uE models

In Scenario 2, we observed different patterns of prediction accuracies of the uE and ME models. The ME model had higher prediction accuracies for DM and MCMDS at all locations. On average (across locations), the uE model had 2 percent better predictive power for VIGOR and 1 percent for RTNO while the ME model had 32 percent better predictive power for DM, 24 percent for MCMDS, 5 percent for FYLD and lastly 4 percent for MCGM. The ME models had 12 percent higher predictive power than the uE model for all traits and locations in the model.

## Discussion

### Scenario 1: MT vs uT model

Few comparisons between MT and uT models have previously been reported in simulation studies and also real data sets [57-59]. Guo et al., 2014 [59] and Calus et al., 2011 [58] in their studies with simulated data sets reported similar accuracies with small differences between their MT and uT models where accuracies for the MT models for low heritability traits were slightly higher when genomic correlations between the traits increased. VanRaden et al., 2014 [57] in Table 10 of their paper (using Holstein and Jersey breed datasets from the US Dairy National evaluation program) also reported similar accuracies with small differences between their MT and uT models for all the traits in their analysis. In several traits, their uT model accuracies were slightly higher than those of their MT model. Accuracies from the MT model will not be clearly better than those from the uT model for traits with high heritability and for traits whose complete phenotypic data are available [59]. Improvement in prediction accuracies for the MT model is accrued mostly for low heritability traits analysed jointly with high heritability traits that have medium to high genomic correlations and low residual correlations [58, 59]. Our results were consistent with other studies [58,59] where our MT model had higher accuracies for most traits and locations in our analysis as a result of joint analysis of low heritability traits with other traits of higher heritabilities. We also observed from pooling data that more efforts on a trial design that replicates lines across multiple locations may improve the predictive power of these models. For parental selections in specific locations, we recommend the use of MT models. However, for varietal selections where non-additive genomic effects are useful, further analysis will have to be performed using data from fully replicated trials at multiple locations (termed uniform yield trials in cassava breeding) to inform this decision. Further studies on the selection gains based on these models are also recommended to facilitate a more accurate decision.

### Scenario 2: ME vs uE model

Again few comparisons of the ME and uE models have been done in plant breeding literature. However, Burgueno et al., 2012 [35] conducted extensive modeling for multi-environment trials using pedigree and genomic markers and incorporated many covariance structures including diagonal, factor analytic (FA), identity and unstructured covariances for both the genomic and error components in their models. They observed higher prediction accuracies for their genomic ME model with a diagonal genomic covariance structure and a diagonal error covariance structure (ME_D-D_) compared to their genomic ME model with a FA genomic covariance structure and diagonal error covariance (ME_FA-D_) for most of the locations in their analysis based on their cross-validation scheme (CV1) [35]. This ME_D-D_ can be likened to our one-step uE model which combined data for a trait measured at multiple locations together and assumes same genomic and error variances for this trait at all locations analyzed and treating multiple location data as just replicates. Our results were in line with this study for the traits VIGOR and RTNO at all our locations where the uE model had higher prediction accuracies than the ME models and differed from this study for the traits DM and MCGM at all locations, FYLD at Ubiaja and Ibadan where the ME models had higher prediction accuracies than those of the uE model. Interestingly on average across locations, the ME models had better predictive power for the most valuable traits in cassava (a combination of DM and FYLD; so called dry yield). Like in other crops, there is need to genetically improve these production (yield) traits in order to drive the economies of the regions that rely on cassava production. Genetic improvement can be made on these cassava target traits using a genetic evaluation system based on multivariate mixed models. The ME model exploits the positive genomic and residual correlations captured in the their *G* and *R* matrices for prediction. The differences between the prediction accuracies of these models (ME and uE) were mainly due to the the estimation of these covariances since their genomic variances were very similar. In addition, the genomic covariances for the ME models are a reflection of GxE interactions of the trait of interest.

Given these genomic covariances from the ME model’s G matrix, we observed differing degrees of GxE ( observing the genomic correlations which were less than 0.9) at the locations Ubiaja, Mokwa and Ibadan (Table 3B). Interestingly, the performance of clones for the traits VIGOR, DM and CMD were most similar between Ubiaja and Ibadan while for the production traits RTNO and FYLD clonal performances were most similar between the locations Ubiaja and Mokwa. Lastly for the performance of clones for MCGM, Ibadan and Mokwa were most similar. A very naive implication of this is that there are no defined clusters for regional breeding by trait and regions as genomic correlations from the ME model partitioned these traits into different categories with different location pairs (Table 3B). From these estimates, improvement for RTNO and FYLD at Ubiaja may translate to about 50 percent stability for these traits at Mokwa but at Ibadan these may translate to about 35 percent. This makes a case for decentralized breeding especially for these yield component traits. These also revealed that cassava DM and MCMDS were highly repeatable across the locations in our study suggesting that more progress can easily be made from selection on these traits and that genotypes selected for these traits will perform appreciably similar across the locations in our analysis and other very closely related locations not in our analysis. However for RTNO, FYLD, VIGOR and MCGM, their GxE profiles reveal that the environment exacts more influence on these traits making their improvement more challenging. Breeding for good varieties which combine these core traits may be targeted towards regional or environment clusters with specific genotypes selected for these clusters. We could also obtain from the GEBVs of the ME model fits a typical reflection of how the clones in our model interacted with the different locations analyzed trait by trait. However, ME GEBVs lack information from trait-trait covariances which has been a difficult hurdle to cross when selecting for multiple traits simultaneously using MET data.

**Table 3A.**
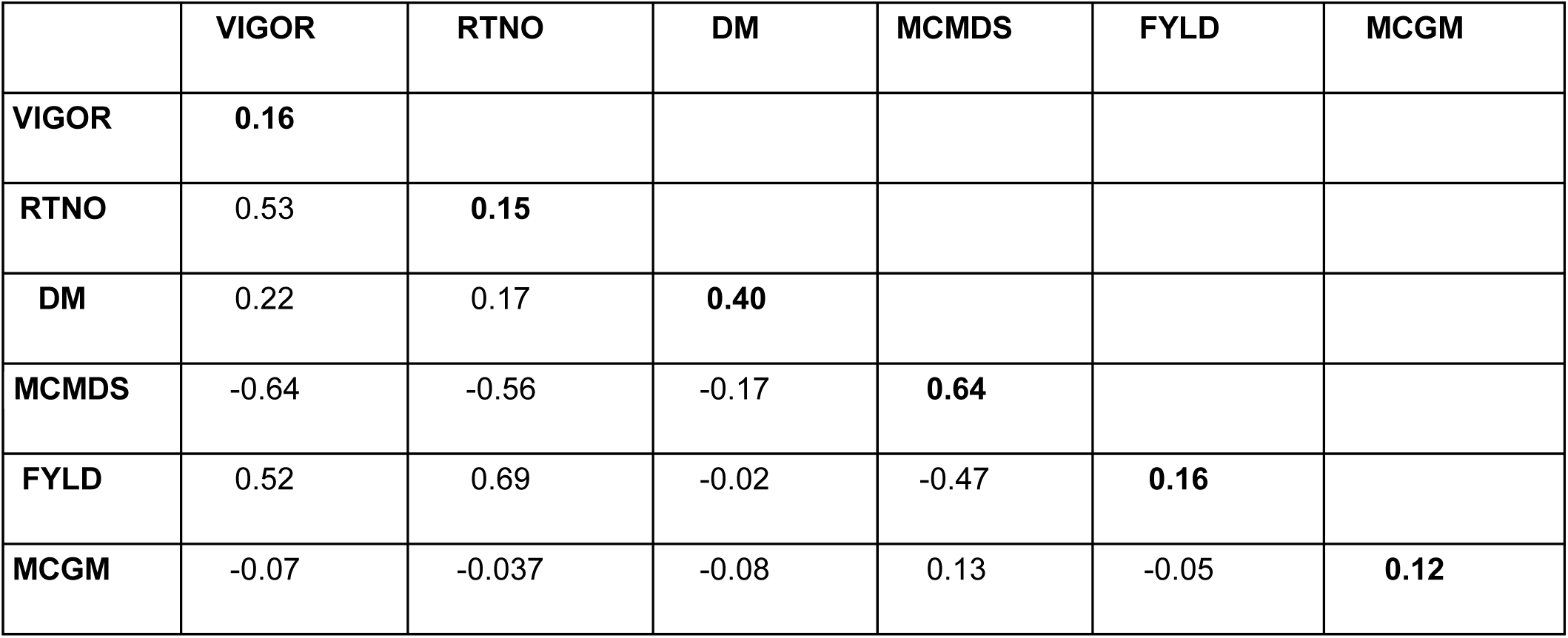
Genetic correlations and heritabilities for analyzed traits. Table 3A shows plot-based heritabilities for six cassava traits on diagonals respectively, genetic correlations from the MT model as lower triangular values.

**Table 3B.** Genetic correlations from the multi-environment analysis. Table 3B show genetic correlations from the ME model for six core cassava traits.

| **VIGOR** |  | **Ubiaja** | **Mokwa** |
| --- | --- | --- | --- |
|  | **Ubiaja** |  |  |
|  | **Mokwa** | 0.39 |  |
|  | **Ibadan** | 0.66 | 0.21 |
| **RTNO** |  |  |  |
|  | **Ubiaja** |  |  |
|  | **Mokwa** | 0.54 |  |
|  | **Ibadan** | 0.36 | 0.38 |
| **DM** |  |  |  |
|  | **Ubiaja** |  |  |
|  | **Mokwa** | 0.57 |  |
|  | **Ibadan** | 0.81 | 0.77 |
| **MCMDS** |  |  |  |
|  | **Ubiaja** |  |  |
|  | **Mokwa** | 0.8 |  |
|  | **Ibadan** | 0.87 | 0.68 |
| **FYLD** |  |  |  |
|  | **Ubiaja** |  |  |
|  | **Mokwa** | 0.52 |  |
|  | **Ibadan** | 0.31 | 0.33 |
| **MCGM** |  |  |  |
|  | **Ubiaja** |  |  |
|  | **Mokwa** | 0.34 |  |
|  | **Ibadan** | 0.24 | 0.53 |

### Parameter estimates and implications for cassava breeding

The estimates of genomic correlations and heritabilities shown in Table 3A have interesting implications for cassava genetic improvement. Most of the target cassava traits (VIGOR, RTNO, FYLD and MCGM) have low heritabilities and hence should benefit from joint analysis with traits of higher heritabilities (DM and MCMDS) in an MT model. The genomic correlations for the core production traits including DM, RTNO and FYLD indicates a promising horizon for improvement. RTNO and FYLD have high positive genomic correlation (0.69) as well as a medium correlation for RTNO and DM (0.17) while that for FYLD and DM is rather weak or almost neutral (-0.02). This indicates that these traits can be improved concurrently with efforts towards fully replicated trials and more accurate parental selections given the low heritabilities for FYLD and RTNO. VIGOR can also be improved concurrently with these production traits as it is also positively correlated with these (Table 3A). The disease trait (MCMDS) shows strong negative genomic correlations with VIGOR and the production traits which raises legitimate concerns towards cassava breeding in Africa where the cassava mosaic disease (CMD) pressure is high. Good progress have been made to fix genes for CMD resistance [64] but a good question to answer is whether this is bringing about negative results and limitations for other traits especially the production traits. This leads to the effort for searching for more quantitative resistance for the diseases CMD and the pest CGM which may better allow for improvement of the traits VIGOR, RTNO, DM and FYLD. Here, the merit index from MT breeding values will offer a good alternative towards a balanced selection goal because it takes into account the genomic correlations of these traits.

## Conclusion

The effectiveness and success of a breeding program is generally evaluated by its impact in its locale mainly on its ability to provide highly adapted and productive varieties to the farming community it serves. This is a huge task to a breeder who will try to fulfil this mandate by making subtle and accurate evaluations of the performance of all the genotypes in his breeding program and subsequently making critical selection decisions based on these evaluations. In our case at the Cassava breeding program at IITA, we are scouting for highly predictive models for use in accurate evaluations of the performance of all our genotypes and hence efficient implementation of GS in our program. In this paper, we have compared methods utilized for this process (in Scenario 1) and compared the efficiencies of these models using prediction accuracies, an indispensable parameter for prediction of selection gains. We also compared models accounting for GxE for genotype evaluations at multiple environments in Scenario 2. We have done these using sixteen years data measured on core or target cassava traits. Genetic evaluation which consists of accurate estimation of breeding values for all individuals in a population for the desired traits of interest and subsequently combination of these breeding values in a total merit index is the crux of genetic improvement. However, the models implemented for estimation of breeding values affect the total merit index and subsequently response to selection. We have shown here that there is value in accounting for covariances using multivariate mixed models for METs data. This value will translate into superior total merit index and hence more selection gain when compared to univariate mixed models. We have also shown that the genomic covariance matrix for ME models can show the effects of GxE for all the traits in the analysis thus helping the breeder to make better decisions. More importantly, we have shown that the MT model has on average 32 percent better predictive power than the uT model while offering more accurate breeding or genetic values for use in total merit indices. We therefore recommend the use of multivariate models (MT and ME) for genetic evaluation in the breeding program where ME is used to assess GxE interactions and MT breeding or genetic values used to set up total merit indices for selection purposes.

## Declarations

### Availability of Data

Phenotypic and genotypic data used in this study will be dumped at Cassava Base (https://www.cassavabase.org/) and will be available for download via an ftp link. Availability is guaranteed upon acceptance of this manuscript.

### Author’s contributions

UGO designed, carried out study and drafted manuscript, DA provided statistical assistance and advice, JLJ supervised the study and revised manuscript, IR and PK supervised data generation for this study and revised manuscript. All authors read and approved manuscript.

## Acknowledgements

We acknowledge the Bill & Melinda Gates Foundation and UKaid (Grant 1048542; http://www.gatesfoundation.org) and support from the CGIAR Research Program on Roots, Tubers and Bananas (http://www.rtb.cgiar.org). We give special thanks to A. G. O. Dixon for his development of many of the breeding lines and historical data we analyzed. Thanks also to A. I. Smith and technical teams at IITA for collection of other phenotypic data and to A. Agbona for data curation.

